# The Proteomic Architecture of Schizophrenia Cerebral Organoids Reveals Alterations in GWAS and Neuronal Development Factors

**DOI:** 10.1101/2021.08.11.455952

**Authors:** Michael Notaras, Aiman Lodhi, Haoyun Fang, David Greening, Dilek Colak

## Abstract

Schizophrenia (Scz) is a brain disorder that has a typical onset in early adulthood but otherwise maintains unknown disease origins. Unfortunately, little progress has been made in understanding the molecular mechanisms underlying neurodevelopment of Scz due to ethical and technical limitations in accessing developing human brain tissue. To overcome this challenge, we have previously utilized patient-derived Induced Pluripotent Stem Cells (iPSCs) to generate self-developing, self-maturating, and self-organizing 3D brain-like tissue known as cerebral organoids. As a continuation of this prior work [1], here we provide a molecular architectural map of the developing Scz organoid proteome. Utilizing iPSCs from *n* = 25 human donors (*n* = 8 healthy Ctrl donors, and *n* = 17 Scz patients), we generated 3D human cerebral organoids, employed 16-plex isobaric sample-barcoding chemistry, and simultaneously subjected samples to comprehensive high-throughput liquid-chromatography/mass-spectrometry (LC/MS) quantitative proteomics. Of 3,705 proteins identified by high-throughput proteomic profiling, we identified that just ~2.62% of the organoid global proteomic landscape was differentially regulated in Scz organoids. In sum, just 43 proteins were up-regulated and 54 were down-regulated in Scz patient-derived organoids. Notably, a range of neuronal factors were depleted in Scz organoids (e.g., MAP2, TUBB3, SV2A, GAP43, CRABP1, NCAM1 etc.). Based on global enrichment analysis, alterations in key pathways that regulate nervous system development (e.g., axonogenesis, axon development, axon guidance, morphogenesis pathways regulating neuronal differentiation, as well as substantia nigra development) were perturbed in Scz patient-derived organoids. We also identified prominent alterations in two novel GWAS factors, Pleiotrophin (PTN) and Podocalyxin (PODXL), in Scz organoids. In sum, this work serves as both a report and a resource whereby researchers can leverage human-derived neurodevelopmental data from Scz patients, which can be used to mine, compare, contrast, or orthogonally validate novel factors and pathways related to Scz risk identified in datasets from observational clinical studies and other model systems.

## INTRODUCTION

Schizophrenia (Scz) is a debilitating brain disorder that occurs in approximately ~1% of the population. While Scz onset typically occurs in early adulthood, subtle brain changes and symptoms often begin emerging years prior to onset during the so-called “prodromal period” [2, 3]. In spite of this, it has remained unclear when Scz neuropathology actually begins to unfold in the brain [1]. For instance, does Scz neuropathology begin a couple of years prior to onset in adolescence when prodromal features progressively emerge? Or does Scz neuropathology begin much earlier in neurodevelopment at a scale that is not yet resolvable? Following decades of investigation, there is now strong epidemiological evidence that indicates risk of Scz may begin to accumulate during *in utero* brain development [4–7]. This includes data from numerous, independent, large-scale populations [4–7]. Critically, it remains unclear if *in utero* risk factors for later Scz onset, such as maternal immune activation, famine, or hormonal/steroid factors, elicit risk by inducing neurodevelopmental alterations or promoting rates of *de novo* mutation [8]. While the latter can’t be ruled out as a potential etiological contributor, the former hypothesis holds strong merit given the highly-regulated nature of cortical development *in utero* and the fact that innumerous Scz risk factors exhibit known roles in central nervous system development. Indeed, some novel biological intermediaries are starting to be discovered which link *in utero* environmental risk factors to potential genetic factors, alterations, and/or vulnerabilities [9]. However, resolving these neurodevelopmental hypotheses of Scz has been difficult. Critically, ethical and technical constraints in accessing human primary brain tissue have arrested progress in delineating the neurodevelopmental trajectory of Scz. These ethical and technical limitations are further compounded by our inability to identify prospective cases of Scz, which has further sequestered our understanding of neurodevelopmental mechanisms of psychosis and has caused a rift between the known epidemiology and the presumed neurobiology of Scz. For instance, in the largest GWAS conducted to date a total of 108 loci of risk were identified – yet, many of these loci (e.g. PTN or PODXL) had unknown disease relevance as well as ambiguously defined neurobiology. Without a means to dissect these factors in human-derived tissue, it is possible that identifying the molecular mediators underlying the ontogeny of disease onset in Scz may continue to be protracted.

Recently, we attempted to overcome these technical and ethical limitations to model early neuropathological features of Scz within human-derived tissue. Namely, we modeled the neurodevelopmental pathology of Scz by harnessing human induced pluripotent stem cells (iPSCs) from healthy adults (Ctrls) and idiopathic Scz patients to generate 3D brain-like tissue known as “cerebral organoids” [1]. Cerebral organoids allow human-specific mechanisms of neural development to be studied while capturing the entirety of the molecular-genetic background of patients. This is a particularly useful model system with respect to “black box” diseases such as Scz, whose neurodevelopmental origins have remained unclear, as it allows spontaneously-emerging neural tissue that is self-organizing and self-maturing tissue to be generated from human donors. Thus, 3D stem cell derived methodologies provide access to a limitless supply of human-derived tissue which can be used to dissect complex diseases defined by “*daunting polygenicity*” [10] under controlled laboratory conditions [11]. Cerebral organoids mimic trimester 1 of early brain development and putatively recapitulate the epigenetic [12], transcriptomic [13, 14], and proteomic [1, 11] architecture that is expected of the developing mammalian brain. This also includes the recapitulation of cortical cell-type diversity and cellular events such as migration [15] and evolutionary mechanisms that support neocortical neurogenesis [16]. Because of this, cerebral organoids have already been used to model prenatal drug/narcotic effects [11], microcephaly [17], macrocephaly [18], Zika virus [19, 20], features of autism [21–23], microdeletion syndromes [24] including 22q11 deletion syndrome [25], hypoxic injury [26], and novel neuropathology of Scz [1, 27–31]. In the case of the latter, Scz-related organoid models have revealed a range of novel phenotypes that may be associated with early neurodevelopmental alterations. This includes diminished responses to electrophysiological stimulation and depolarization [27], alterations in growth factor pathways (e.g. FGFR1 [28] and neurotrophic growth factors and their receptors in Scz progenitors and neurons [1]), immune-related alterations (e.g. TNFα [29] and IFITM3 as well as IL6ST in Scz neurons [1]), potential developmental effects in excitation and inhibition [30], and DISC1 effects upon neurodevelopment [31, 32]. Recently, we added to this developing literature by being the first to discover that Scz neuropathology is encoded on a cell-by-cell basis and is defined by multiple novel mechanisms in Scz patient-derived organoids [1]. However, we have also predicted that further mechanisms related to neurodevelopment of Scz remain to be discovered [1], thus requiring deeper analysis in larger samples and populations.

Here we sought to expand our existing knowledge of Scz by providing a deep, unbiased, analysis of molecular factors regulating central nervous system development in human-derived 3D tissue. To do this, we generated cerebral organoids from a relatively large pool of human donors (*n* = 25; *n* = 8 Ctrl donors and *n* = 17 Scz donors) and adapted cutting-edge isobaric barcoding chemistry so that samples could be condensed and analytically deconstructed simultaneously via liquid-chromatography/mass-spectrometry (LC/MS). This yielded a large dataset that we have made freely available for other human, mouse, and cellular researchers to analyze. Notably, here we emphasize large-scale changes identified in this dataset, which included a broad reduction in neuronal molecules important for neural cell-type identity and development as well as metabolic and novel GWAS factors. This work and dataset may thus provide insight for other researchers and labs that have an interest in biological data from human-derived 3D stem cell systems but otherwise employ or use other model systems.

## RESULTS

To study the molecular architecture of developing human brain-like tissue, we generated 3D cerebral organoids from human iPSC donors banked by the NIMH. In sum, biologics from *n* = 25 human donors were sampled comprising *n* = 8 healthy Ctrls and *n* = 17 Scz patients. Briefly, iPSCs from human donors were grown in 2D culture atop vitronectin-coated plates before being dissociated with Accutase to yield single-cell iPSCs suspensions. Stem cell suspensions were correspondingly cultured into 3D aggregates, known as embryoid bodies, before being subjected to a chemically minimalist neural induction media for up to 7 days *in vitro* (DIV). After exhibiting evidence of neuroepithelial expansions and/or other morphological evidence of neural induction, tissue was impregnated into a matrigel droplet as a scaffold for further tissue expansion. Developing organoids were then maturated under constant agitation atop an orbital shaker. Following this, at approximately 35-40 DIV, organoids from all 25 human donors were sampled for TMT quantitative proteomics. Briefly, this involved dissociating organoids, preparing peptide suspensions (digestion, reduction, and alkylation), barcoding samples with isobaric TMTpro 16-plex chemistry, and then multiplexing samples for simultaneous detection and analysis via nano high-sensitivity proteome profiling (for a simplified schematic of our experimental pipeline, see Fig. 1).

**Figure 1.**
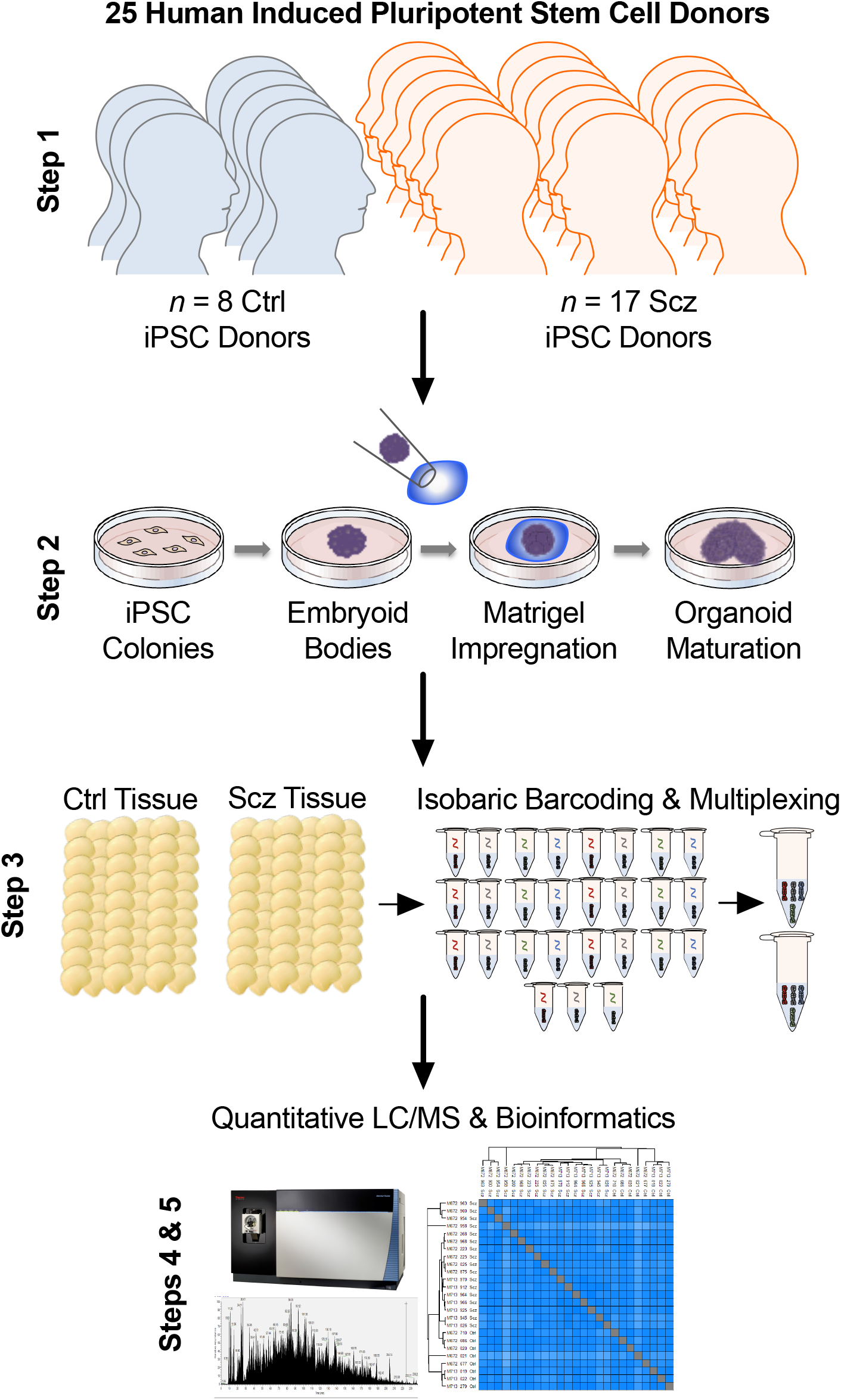
Schematic of cerebral organoid and TMT-LC/MS analytical pipeline. Briefly, 25 distinct human iPSCs were obtained from both healthy Control (Ctrl) donors and Schizophrenia (Scz) patients. Each line represented a biologically unique sample from a specific individual, and lines were predominantly obtained from NIH repositories. Following this, iPSCs were expanded and utilized to generate patient-derived cerebral organoids that mimic the 1^st^ trimester of brain assembly (see Methods, [17, 88] for protocol information, and [1] for our previous application of 3D Scz patient-derived organoids). This process involved dissociating iPSC colonies to generate 3D embryoid body aggregates that could be pushed towards a neural fate via chemically minimalist media cocktails [17, 88]. Following neural induction, organoids were implanted into a matrigel droplet as a scaffold to support tissue expansion and, consistent with our prior study [55], maturated to a primary endpoint of 30DIV. Following this, samples were individually subjected to protein lysis and tryptic-based enzymatic digestion. For proteomic analysis of cerebral organoids, peptides were isobarically barcoded using TMTpro 16-Plex chemistry that allowed samples to be multiplexed for simultaneous analysis of different samples via liquid-chomatography/mass-spectrometry. This allowed up to 15 samples (+1 pool) to be condensed into a single tube for simultaneous detection via liquid-chromatography mass-spectrometry (LC/MS) analysis, resulting in a total of 27 samples (*n* = 25 human donor organoids, +*n* = 2 internal reference pools). Proteomic nano-LC tandem mass spectrometry analysis was performed on a Fusion Lumos to molecularly map the protein composition of *n* = 25 of our iPSC human donor samples. Bioinformatics were subsequently conducted in accordance with the parameters described in our Methods as well as two prior manuscripts that have incorporated LC/MS proteomic analysis of human-derived organoid samples [1, 11].

Analysis of organoid proteomes revealed sufficient peptide coverage for high-confidence quantitative analysis of 3705 proteins (peptide >1; intensity > 0) across all 25 human donor samples. Based on Log2 transformed protein intensities, the Coefficient of Variation (CV) of Scz and Ctrl proteome groups was highly stringent; Median CV for Ctrls was 1.07% and for Scz 1.23%. This provided confidence in both the degree of neural induction achieved between samples, and that organoids were overall of a very similar and thus comparable composition between iPSC donors and within groups.

To gain insight into differences between Scz and Ctrl organoids, we next sough to determine which proteins (based on their expression) differed between these groups. Further analysis revealed the significant differential expression of peptide fragments belonging to 97 proteins in Scz organoids, of which 43 were up-regulated (*p* value < 0.05, Log2FC > 0.05) and 54 were down-regulated (*p* value < 0.05, Log2FC < −0.05). Thus, in sum, ~2.62% of the total organoid proteome was differentially expressed in Scz organoids, with equivalent (~1.16% vs. ~1.46%) proportions of differentially expressed proteins being up- and down-regulated, respectively.

Deeper examination of significantly down-regulated proteins in Scz organoids, sorted by Log2FC values (see Table 1), revealed several important changes. Notably, we detected a depletion of factors that support neuronal development, differentiation, identity and/or function. Down-regulated neuronal development factors in Scz organoids comprised Neuromodulin (GAP43; Log2FC = −1.183, *p* = 0.010), Cellular Retinoic Acid-Binding Protein 1 (CRABP1; Log2FC = −1.018, *p* = 0.016), Neural Cell Adhesion Molecule (NCAM1; Log2FC = −0.854, *p* < 0.014), and expression of the myelin-modulating factor Myelin Expression Factor 2 (MYEF2; Log2FC = −0.537, *p* < 0.001). Likewise, down-regulated expression of several other neuronal factors – involved in both neuronal identity and prototypic function - included Microtubule-Associated Protein 2 (MAP2), Tubulin Beta-3 Chain (TUBB3, or β3), Synaptic Vesicle Glycoprotein 2A (SV2A), among other neuron-specific markers (see Fig. 2). In addition to these changes, we also screened our dataset against novel, yet statistically prominent, Scz GWAS factors identified in the largest population genetic dataset reported to date [33]. One important Scz GWAS factor to emerge from our analysis of down-regulated proteins in Scz organoids was Pleiotrophin (PTN). In our prior work [1], we also detected the differential expression of PTN at both the protein and RNA level in Scz organoids, including in both Scz progenitors and neurons. This better powered analysis therefore replicates this previous finding, and further establishes PTN as a potentially important Scz risk factor during early brain assembly.

**Table 1.**
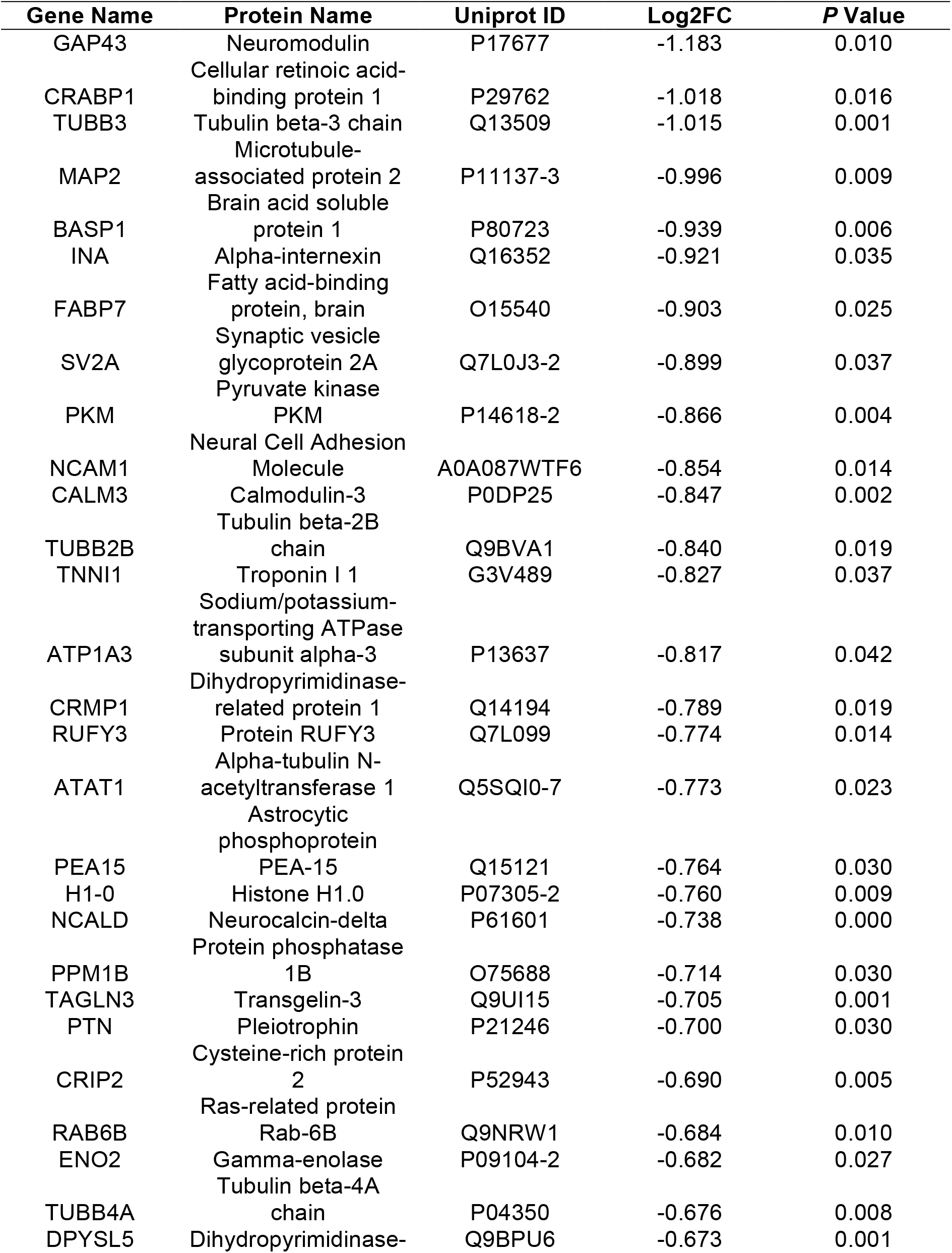
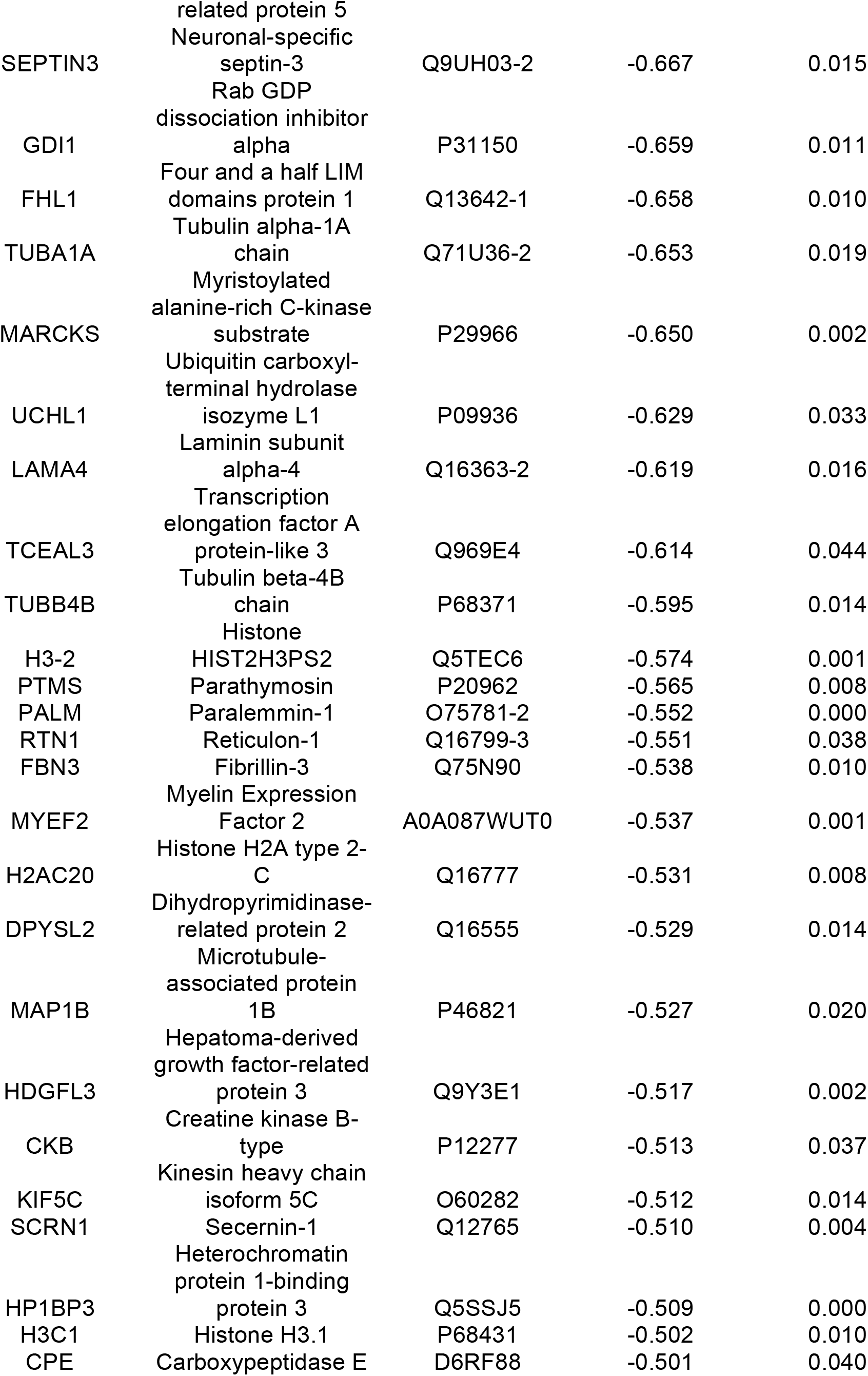
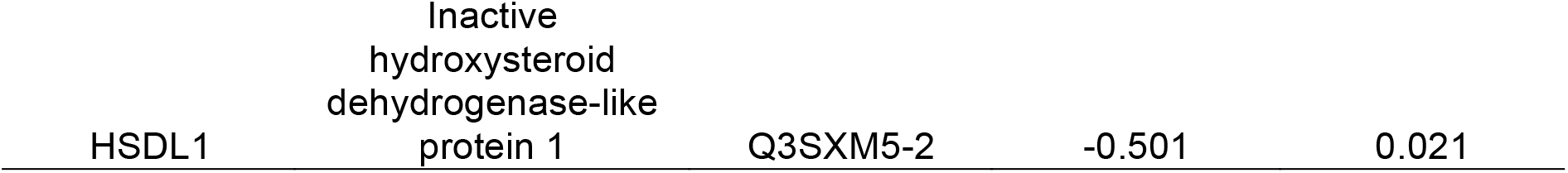
54 Down-Regulated Proteins in Scz Organoids (< −0.5 Log2FC, *p* < 0.05).

**Figure 2.**
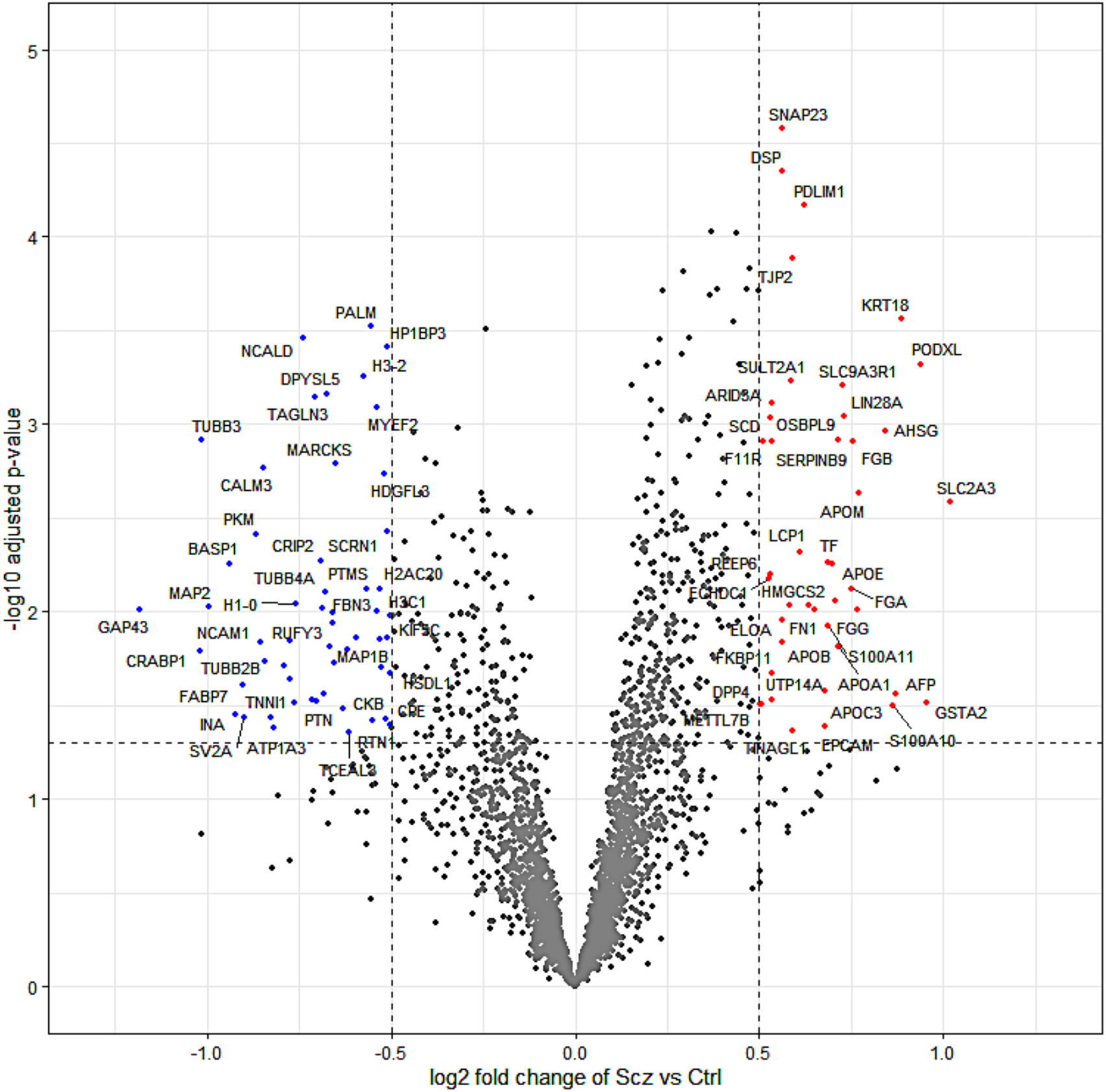
Differential Expression in the Scz Cerebral Organoid Proteome. Principal component analysis of the cerebral organoid proteome indicated data grouping based on phenotype, and protein expression distributions indicated data correlation across all samples. This statistical baseline allowed us to consider the differentially expressed proteins present in Scz patient-derived cerebral organoids, which are shown here as a volcano plot split by log2 fold change and –log10 adjusted *p* values. In sum, ~2.62% of 3705 proteins (peptide >1; intensity > 0) identified exhibited differential expression. Significantly up-regulated proteins that surpassed log2 fold change thresholding are depicted to the right in red (*p* value < 0.05, Log2FC > 0.05), whereas down-regulated proteins (*p* value < 0.05, Log2FC < −0.05) are presented to the left of the plot in blue. Notable Scz GWAS factors (see 108 loci identified in [33]) included the up-regulation of PODXL and down-regulation of PTN, which replicated our previous findings in a smaller cohort [1]. Note also the down-regulation of the neural stem cell proliferation factor CRABP1 [94] as well as canonical neuronal development markers (e.g. NCAM1 [95], NCALD [96], and CPE [79]), neuronal markers (e.g. MAP2, TUBB3, MAP1B), synaptic markers (e.g., SV2A). Conversely, a range of apolipoproteins (APOE, APOA1, APOB, APOC3) were found to be up-regulated in Scz patient-derived cerebral organoids.

Similar to our review of down-regulated proteins, we also identified a number of biologically interesting observations in our up-regulated Scz protein set list (see Table 2). This included up-regulation of numerous fibrinogens (FGG, FGB, FGA; Log2FC = 0.749-0.768, *p* = 0.008-0.010) and apolipoproteins (APOM, APOA1, APOE, APOC3, APOB; Log2FC = 0.562-0.771, *p* = 0.001-0.015). However, one of the most notable up-regulated protein was another Scz GWAS factor [33] that (like PTN) we had also previously identified in our prior Scz patient-derived organoid work [1]; namely, Podocalyxin (PODXL; Log2FC = 0.939, *p* < 0.001). Therefore, similar to our replication of down-regulated PTN expression in Scz organoids, this analysis in a larger pool of patients confirms that PODXL is another high-confidence candidate that may play a role in modulating Scz risk during early brain development.

**Table 2.**
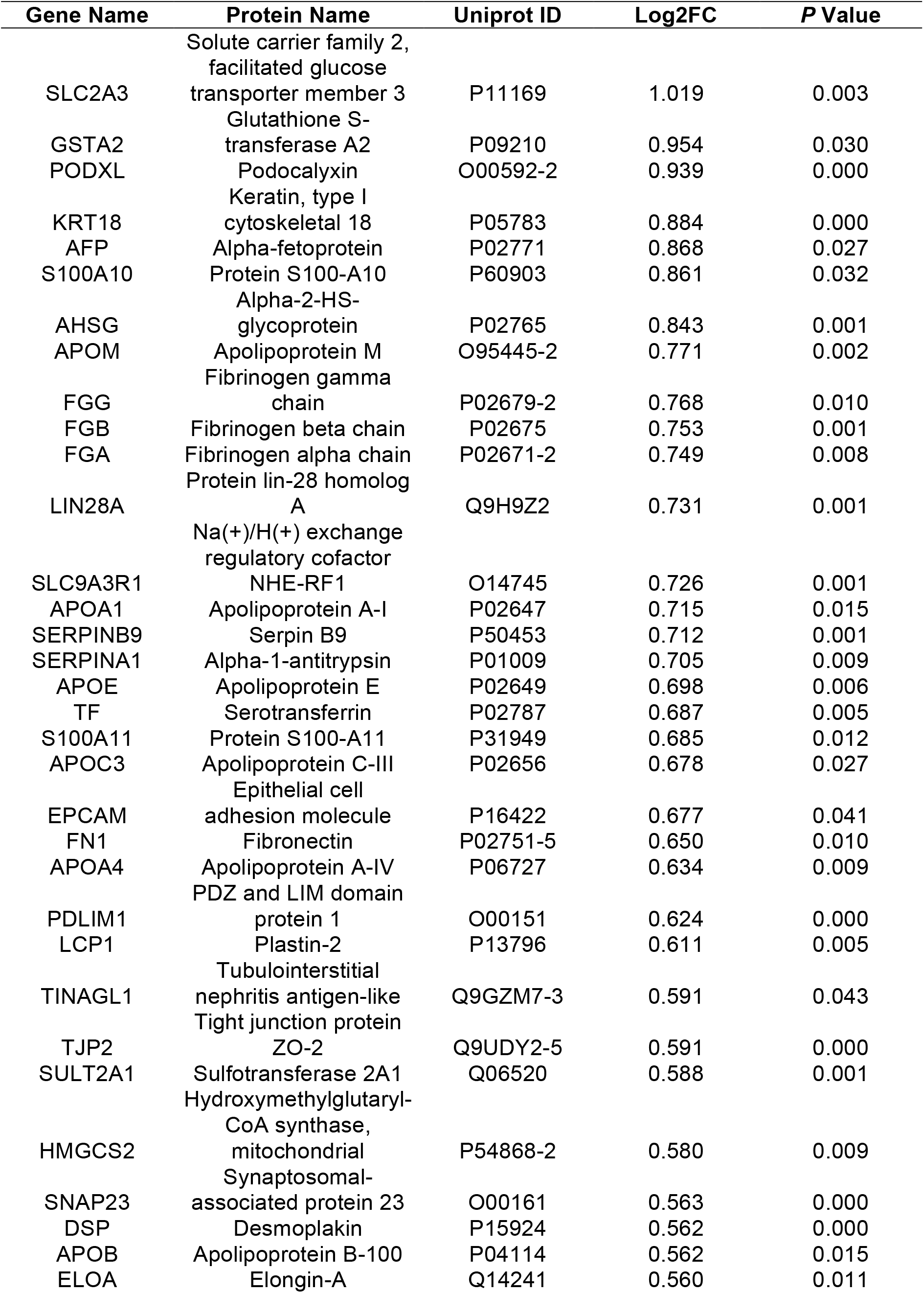
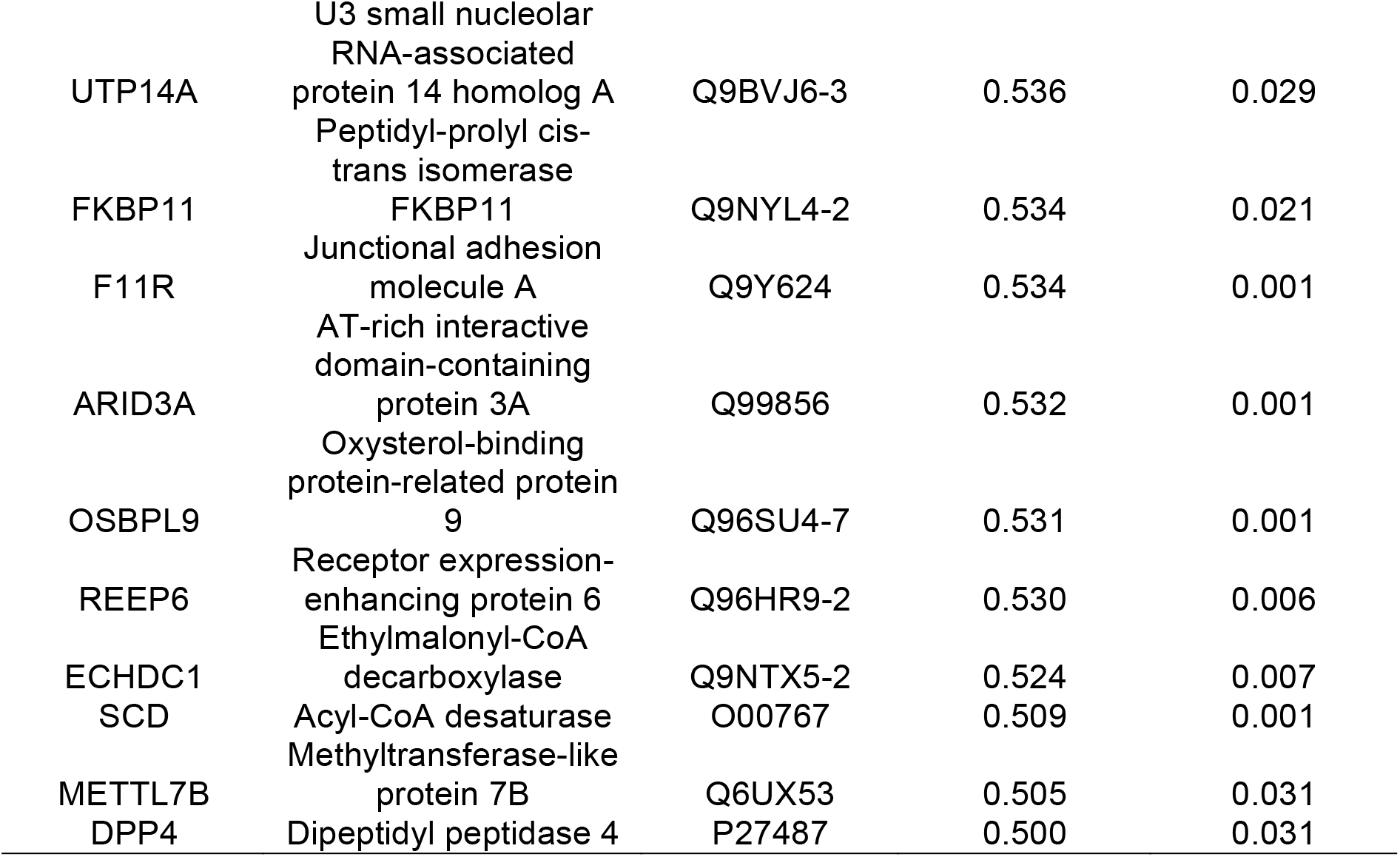
43 Up-Regulated Proteins in Scz Organoids (> 0.5 Log2FC, *p* < 0.05).

We next sought to understand the potential functionality of our differentially expressed protein targets by parsing these factors into pathways, which may also unveil broader changes in regulatory networks underscoring disease-related phenotypes. We principally examined Gene Ontology (GO) pathways, parsed by annotations belonging to biological (Tables 3–4) and molecular (Tables 5–6) function of differentially expressed proteins. We first considered down-regulated GO biological pathways. Down-regulated GO biological pathways essential for normative brain assembly, development, and maturation overwhelmingly defined Scz patient-derived organoids. This included down-regulated expression of factors that map to axonogenesis, axon development, axon guidance, morphogenesis pathways regulating neuronal differentiation, and, broadly speaking, central nervous system development (due the sheer number of pathways involved here, please refer to Table 3 for statistical values). Another interesting down-regulated GO biological process pathway in Scz organoids was specific enrichment for factors regulating substantia nigra development (GO:0021762, adjusted *p* = 0.0182, Neg Log10 = 1.74), which is of interest given that this midbrain region belongs to the basal ganglia which holds broad relevance to Scz neuropathology and its treatment (e.g. dopamine and monoamine hypotheses of Scz development and symptoms). Contrary to down-regulated GO biological pathways, up-regulated pathways in Scz organoids broadly reflected pathways involved in cellular metabolism, chylomicron assembly and remodeling, sterol and steroid pathways, as well as lipoprotein remodeling and metabolism-related pathways (refer to Table 4 for statistical values).

**Table 3.**
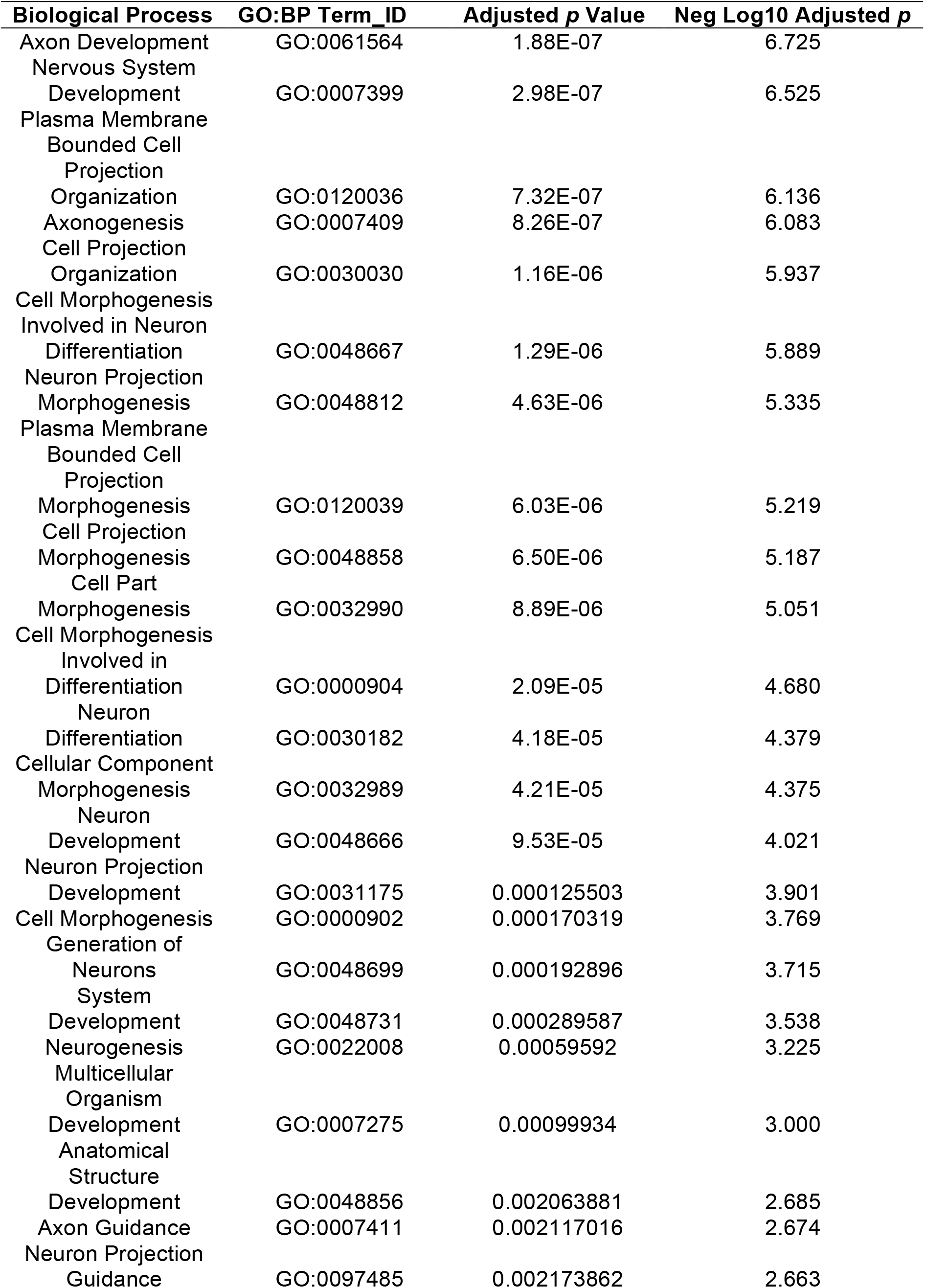
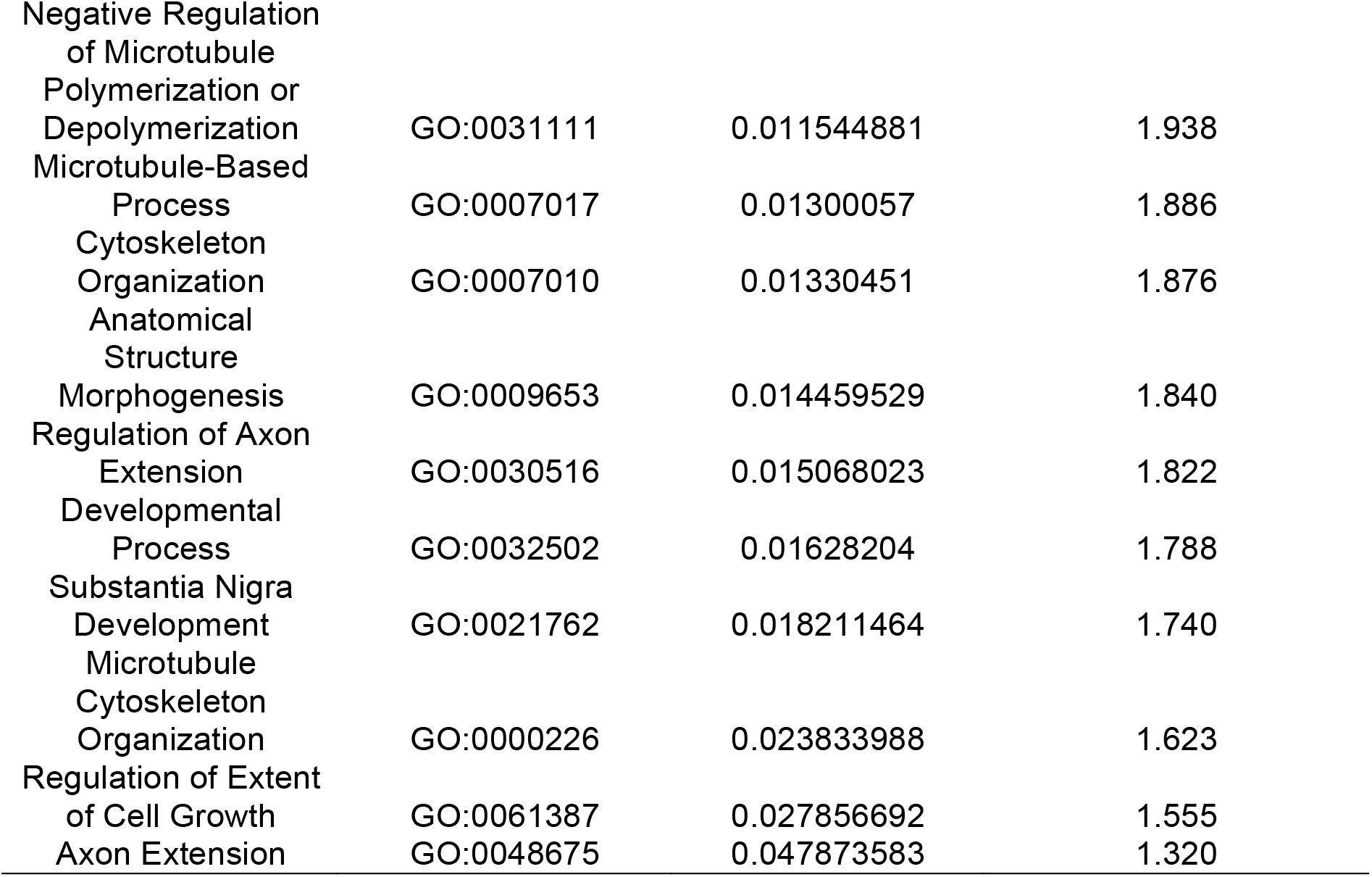
Down-Regulated GO Biological Processes in Scz Organoids (*p* < 0.05).

**Table 4.**
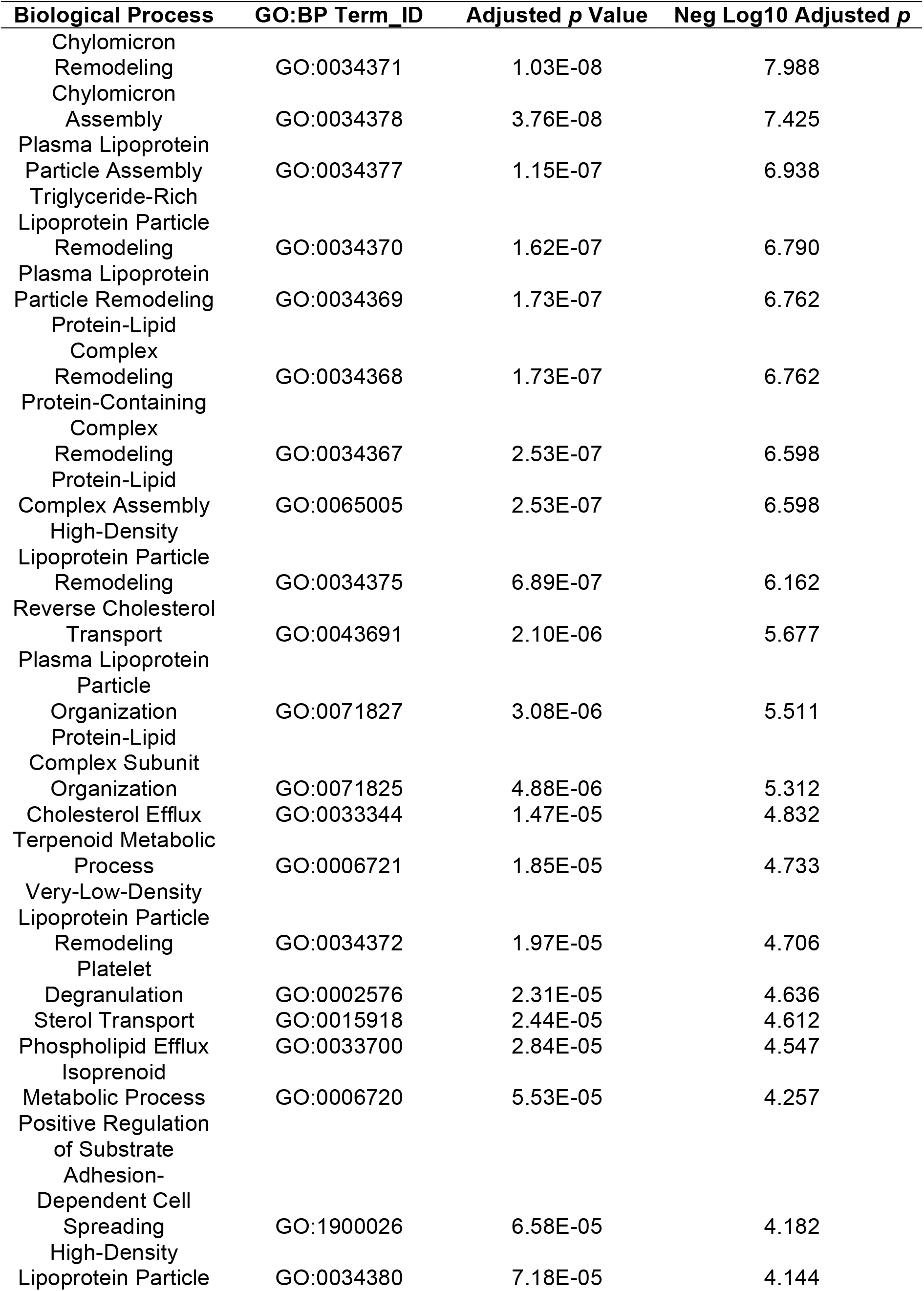
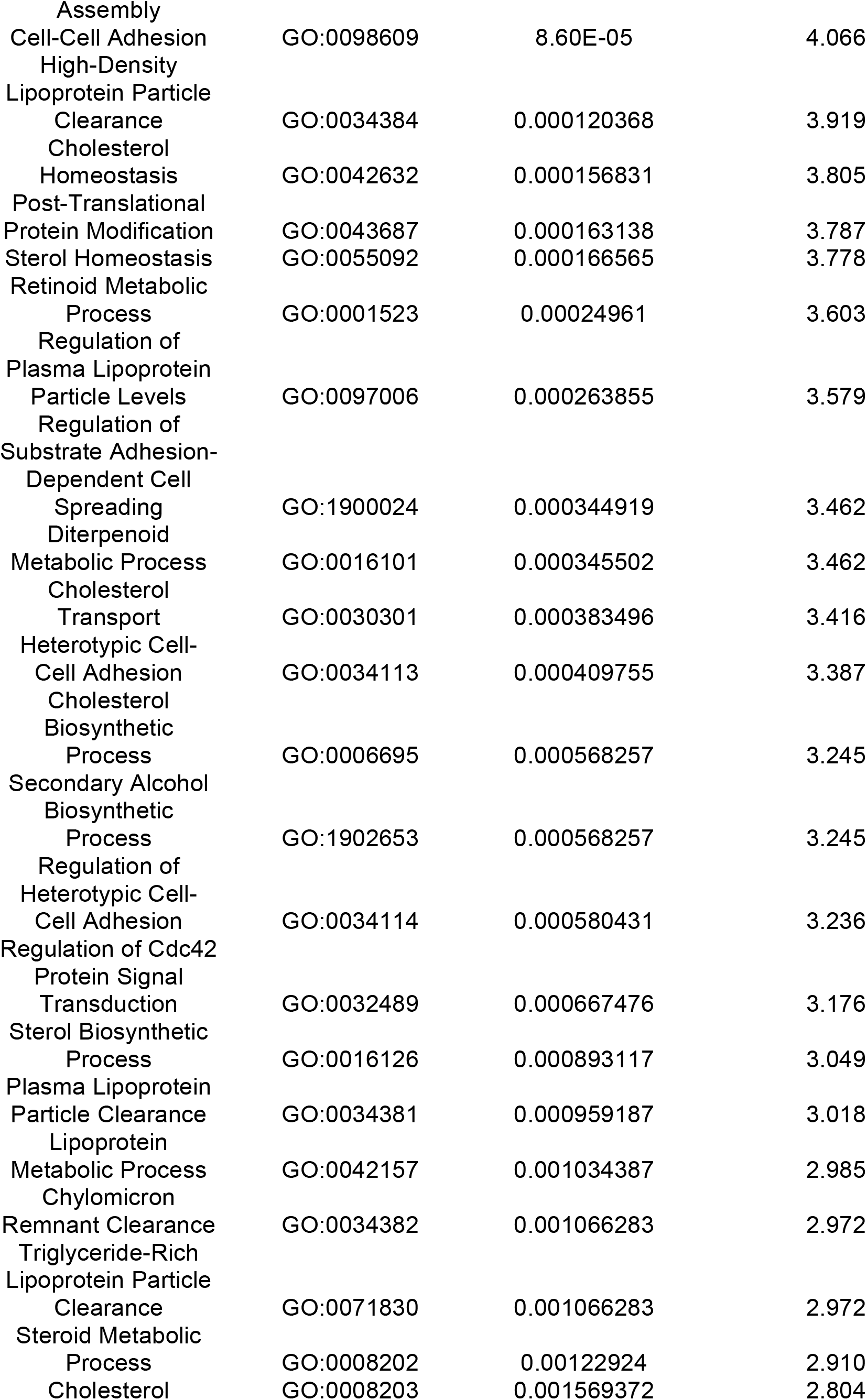
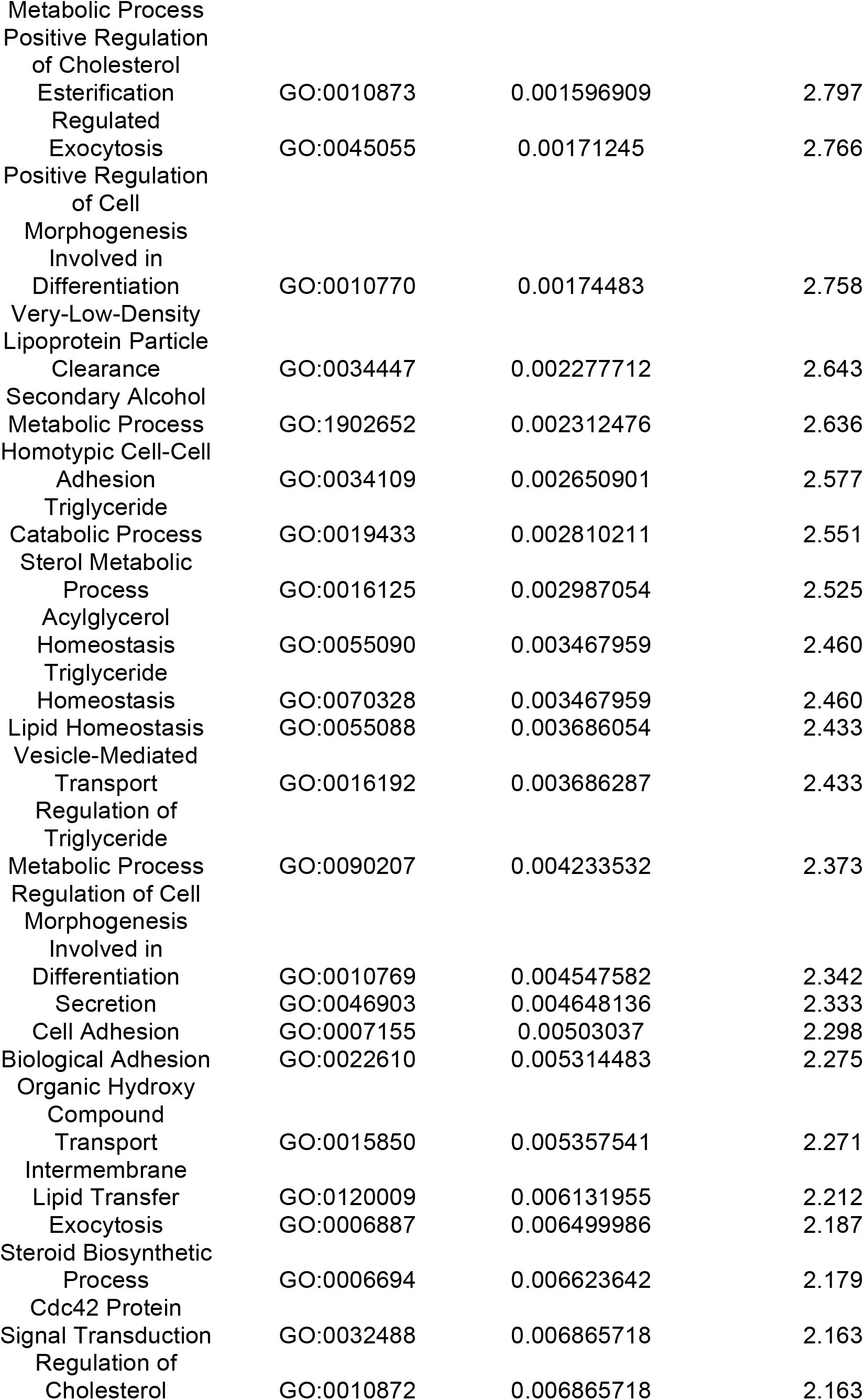
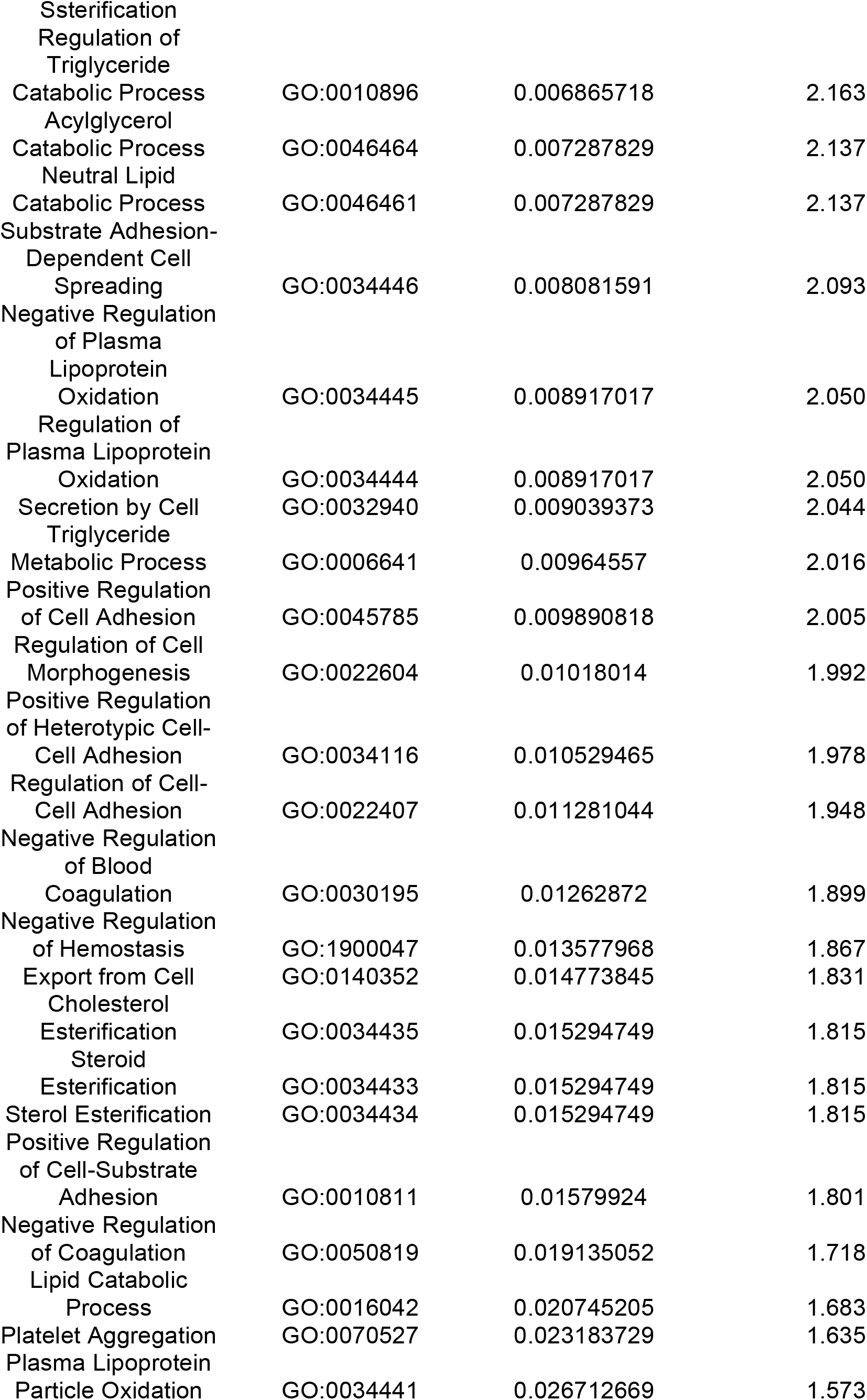
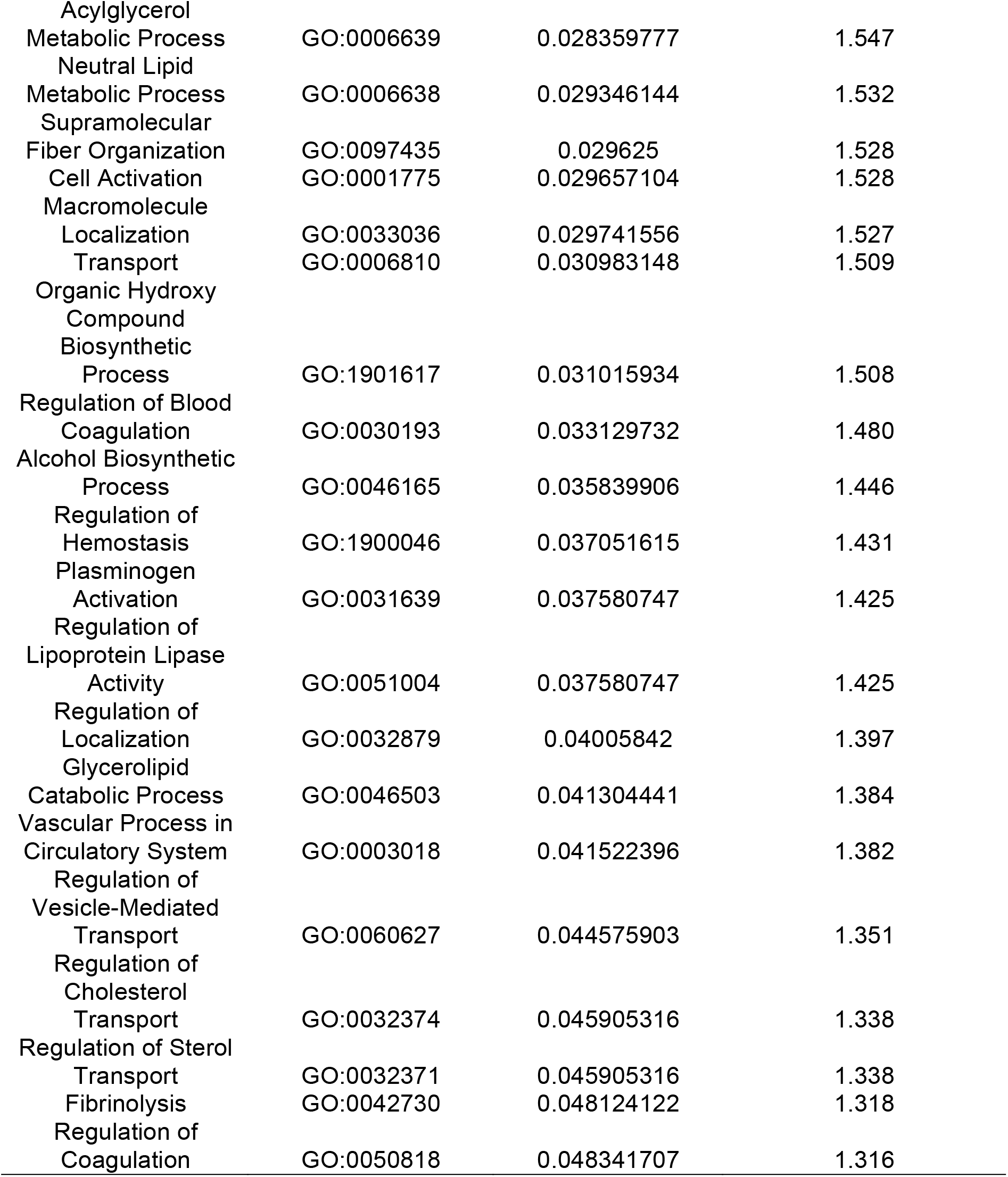
Up-Regulated GO Biological Processes in Scz Organoids (*p* < 0.05).

**Table 5.** Down-Regulated GO Molecular Functions in Scz Organoids (*p* < 0.05).

| **Molecular Function** | **GO:MF Term_ID** | **Adjusted $p$ Value** | **Neg Log10 Adjusted $p$** |
| --- | --- | --- | --- |
| Structural Constituent of Cytoskeleton | GO:0005200 | 0.000173652 | 3.760 |
| Cytoskeletal Protein Binding | GO:0008092 | 0.005488124 | 2.261 |
| GTPase Activity | GO:0003924 | 0.007451524 | 2.128 |
| Nucleoside-Triphosphatase Activity | GO:0017111 | 0.008195516 | 2.086 |
| Pyrophosphatase Activity | GO:0016462 | 0.020438022 | 1.690 |
| Hydrolase Activity, Acting on Acid Anhydrides, in Phosphorus-Containing Anhydrides | GO:0016818 | 0.023962071 | 1.620 |
| Hydrolase Activity, Acting on Acid Anhydrides | GO:0016817 | 0.024306171 | 1.614 |
| Tubulin Binding | GO:0015631 | 0.034081605 | 1.467 |
| GTP Binding | GO:0005525 | 0.034081605 | 1.467 |
| Microtubule Binding | GO:0008017 | 0.043893561 | 1.358 |
| Structural Molecule Activity | GO:0005198 | 0.047624863 | 1.322 |
| Guanyl Ribonucleotide Binding | GO:0032561 | 0.047641356 | 1.322 |
| Guanyl Nucleotide Binding | GO:0019001 | 0.047641356 | 1.322 |

**Table 6.**
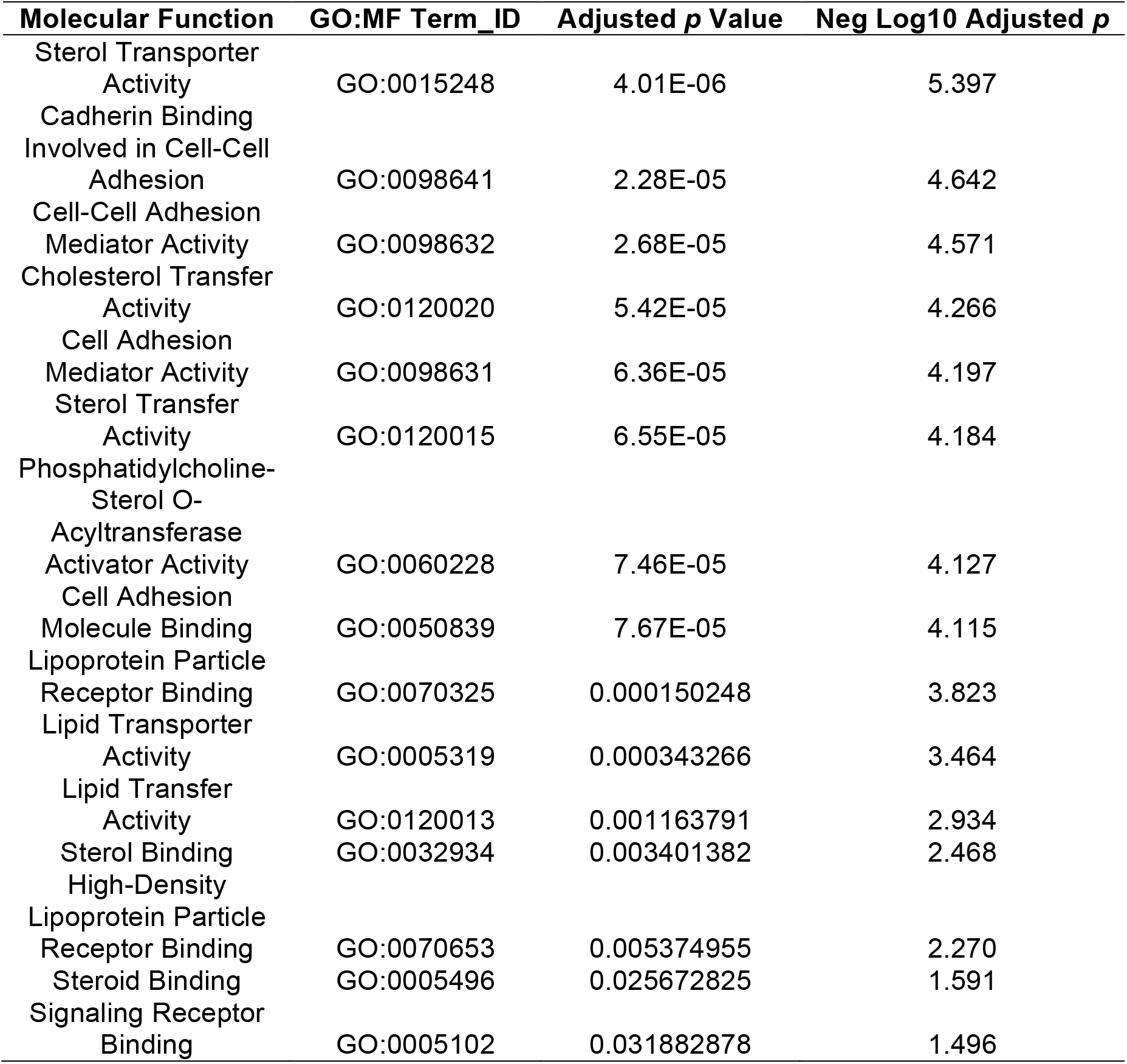
Up-Regulated GO Molecular Functions in Scz Organoids (*p* < 0.05).

Broadly speaking, these changes were also reflected in our analysis of GO pathways annotated for molecular functionality. Specifically, down-regulated GO molecular functions in Scz organoids comprised cytoskeletal structural, binding, and activity, as well as metabolic pathways relevant to neurodevelopment such GTP binding and GTPase activity (see Table 5; also identified in our prior prenatal drug modeling organoid work [11]). Similarly, up-regulated GO molecular function pathways in Scz organoids were typically related to sterol activity, cell adhesion, and lipoprotein binding/transfer/activity (see Table 6). In sum, these data provide additional veracity to the idea that there are metabolic functions underscoring the depletion of neuronal development factors in Scz organoids.

Lastly, we also considered whether Reactome pathways might unveil other novel biology in Scz organoids. Overall, an analysis of down-regulated (Table 7) and up-regulated (Table 8) Reactome pathways in Scz organoids revealed broadly similar pathway enrichment to those identified via GO analysis, with some notable exceptions. First, in our down-regulated Reactome pathway analysis, we noted that there were numerous significant pathways involved in NMDA receptor activation and assembly, ER to Golgi transport, as well as synaptic transmission (see Table 7 for a comprehensive list and statistical values). Contrary to this, and in addition to a convergent detection of lipoprotein-related metabolism pathways, unique Reactome pathways that were up-regulated in Scz organoids comprised post-translational protein phosphorylation, pathways related to MAPK signaling, IGF-related pathways. Overall, these data suggest that ying-and-yang alterations in Scz organoids exist, whereby the disruption of neuronal-development factors and pathways yields enrichment for pathways presumably involved in either compensation or other disease-related neuropathology including phenotypes that have possibly not yet articulated in human-derived tissue (e.g. metabolic changes).

**Table 7.**
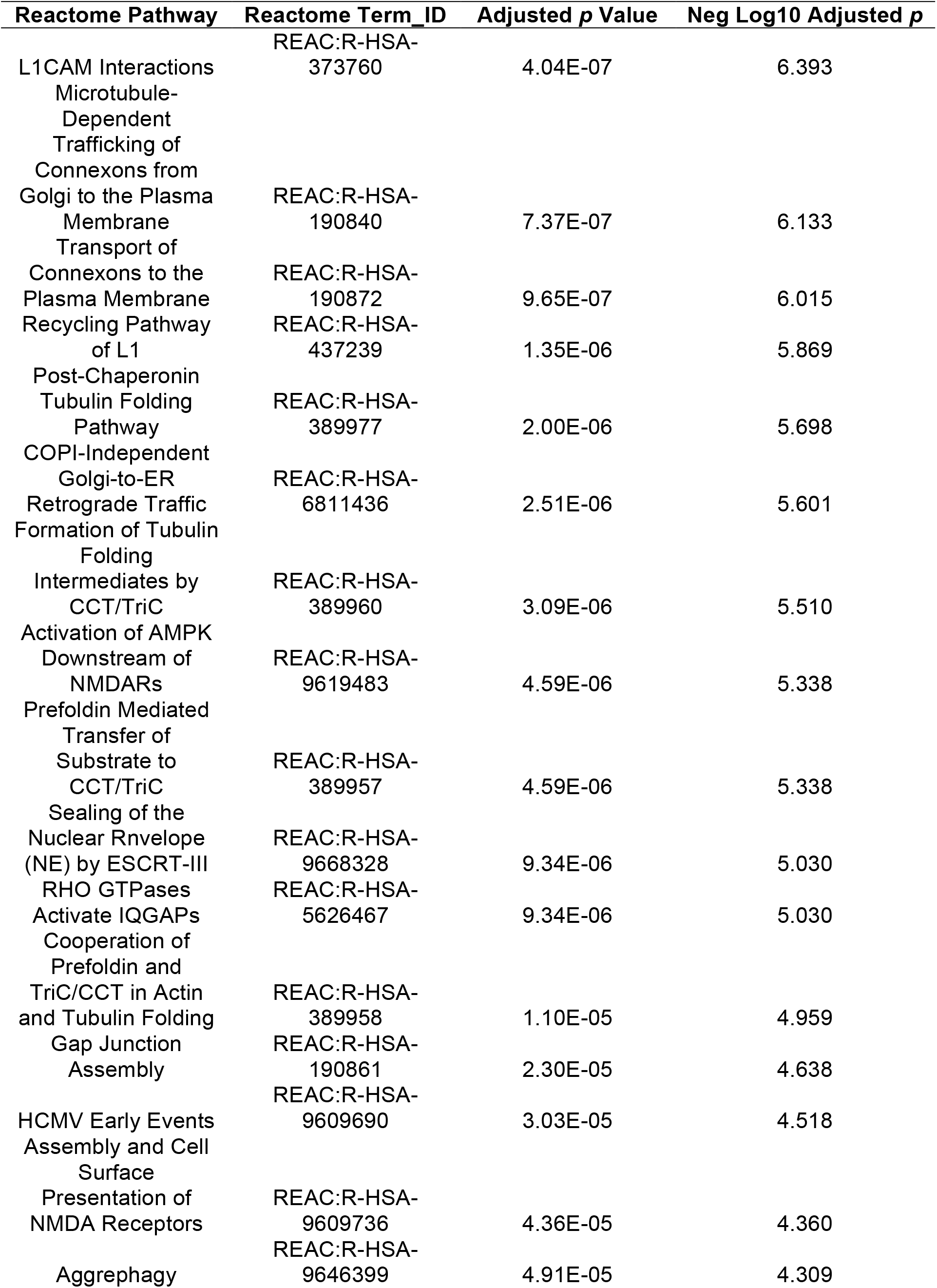
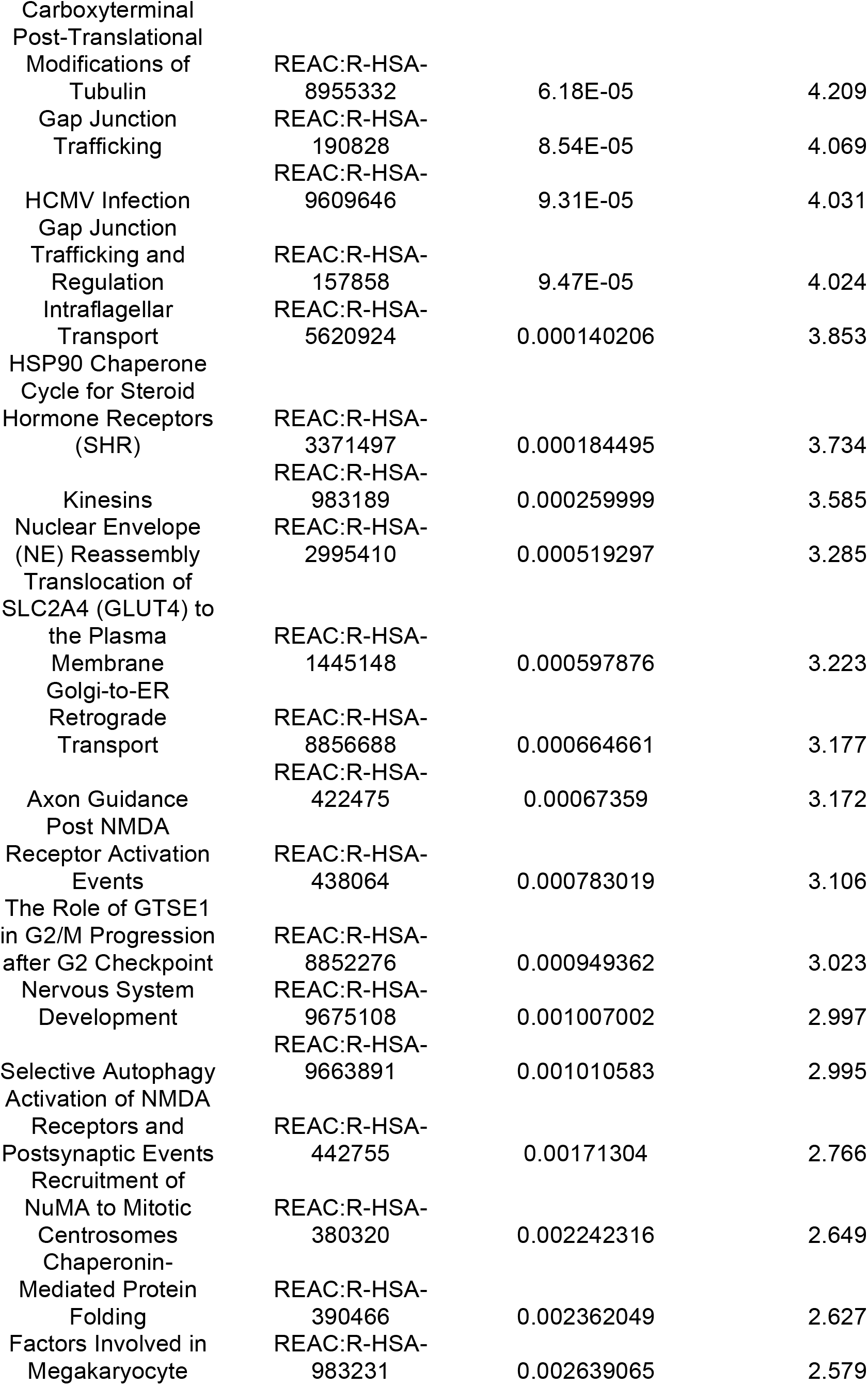
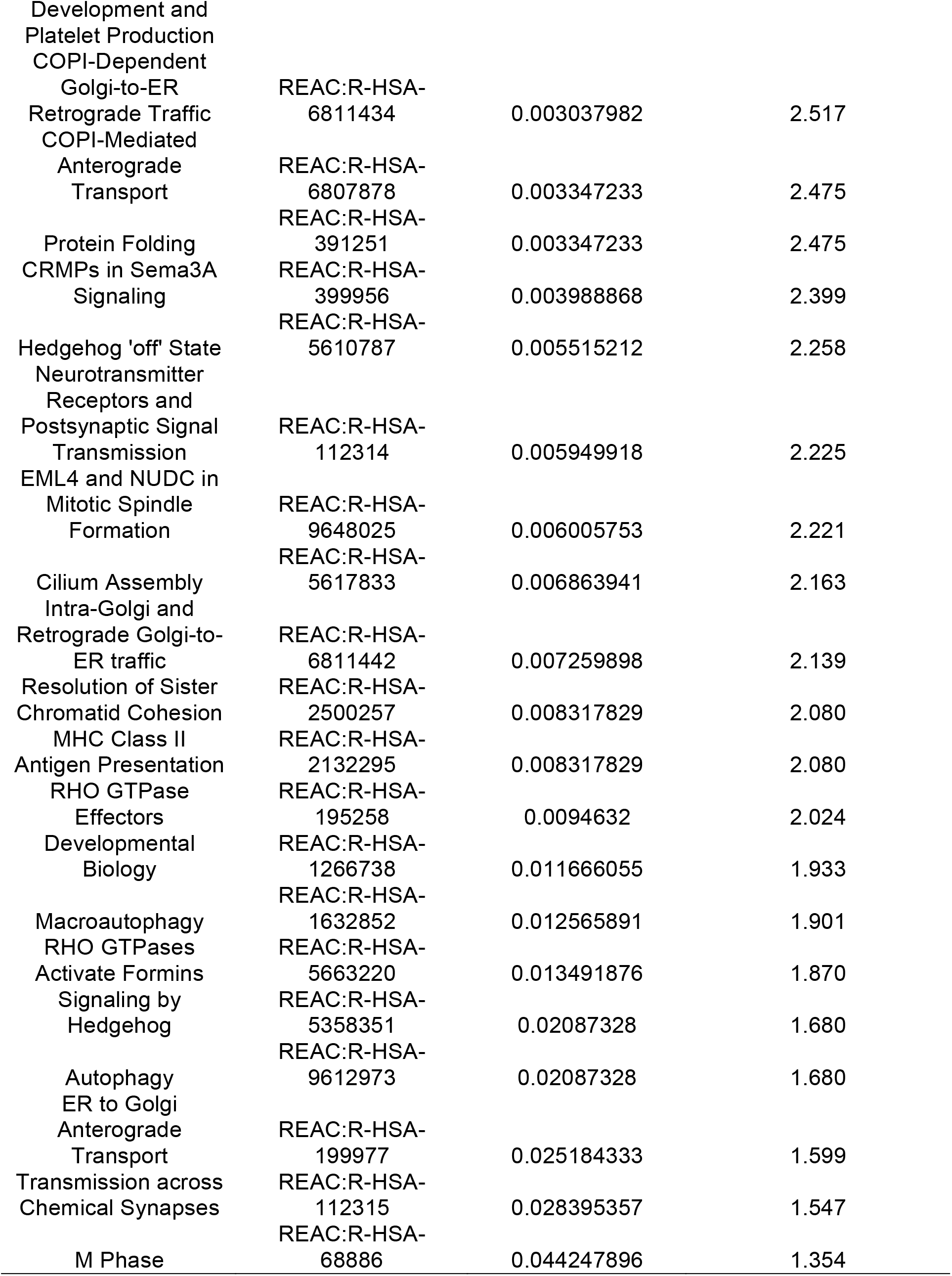
Down-Regulated Reactome Pathways in Scz Organoids (*p* < 0.05).

**Table 8.**
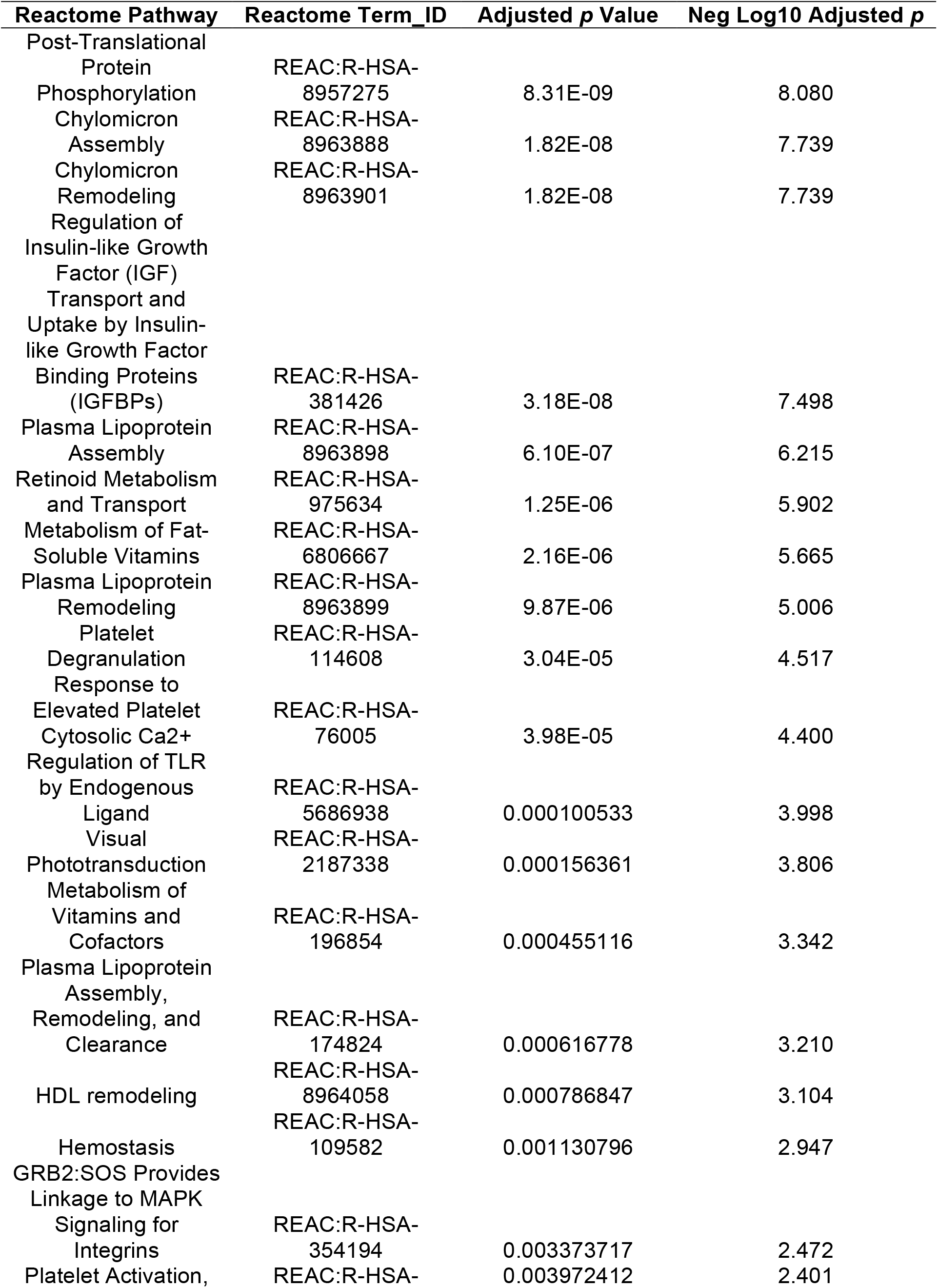
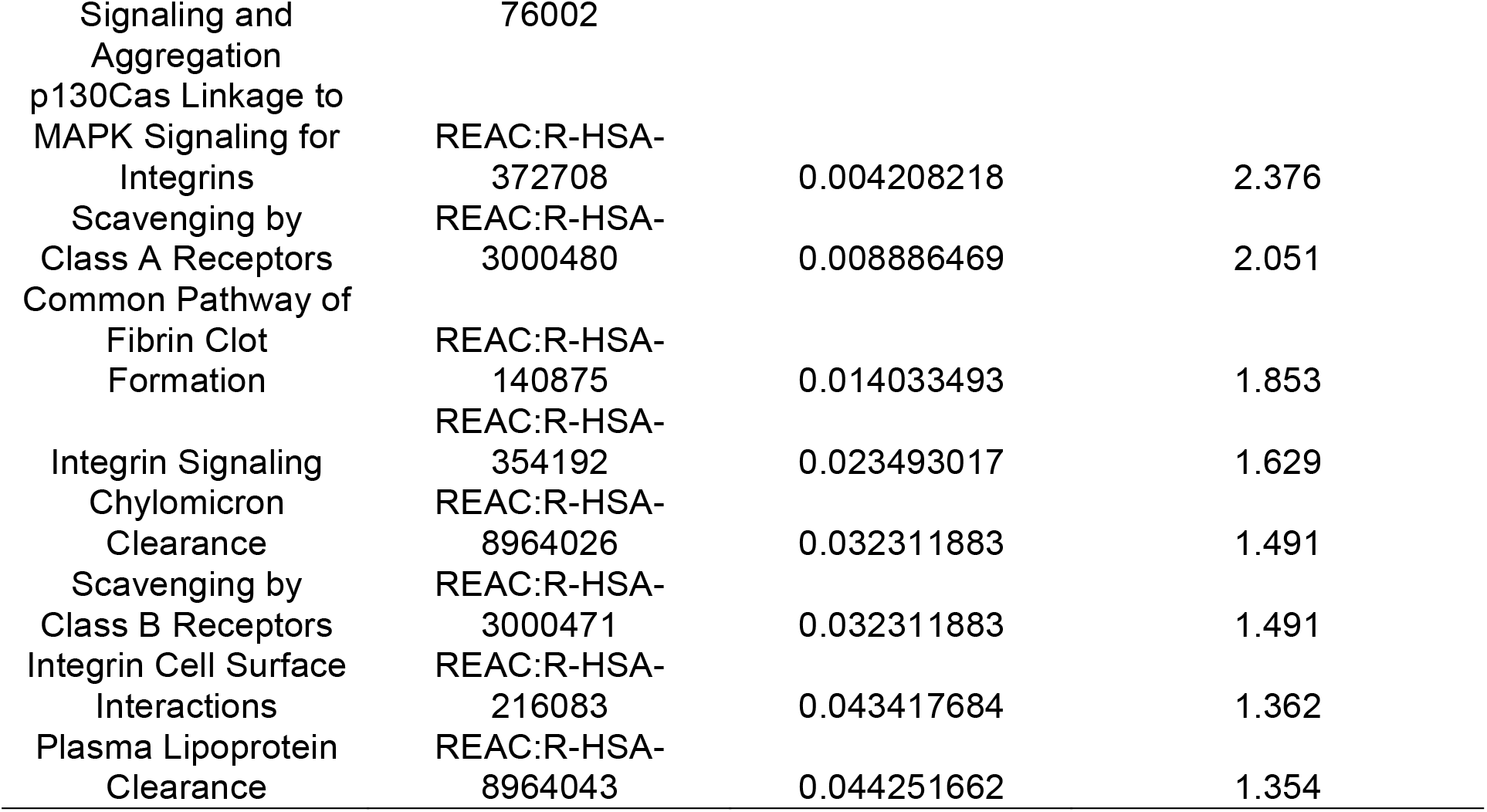
Up-Regulated Reactome Pathways in Scz Organoids (*p* < 0.05).

## DISCUSSION

The aim of the current study was to further our knowledge of Scz by providing a deep, unbiased, analysis of molecular factors regulating central nervous system development in human-derived 3D tissue. To circumvent ethical and technical limitations in being able to access developing neural tissue from Scz patients [11], we generated 3D iPSC-derived cerebral organoids from *n* = 25 human donors (*n* = 8 Ctrl donors and *n* = 17 Scz donors). This approach allowed us to generate a theoretically limitless supply of self-regulating 3D neural tissue that recapitulated hallmark features of early brain assembly and corticogenesis [34, 35]. Samples were correspondingly subjected to cutting-edge isobaric barcoding chemistry that allowed up to 15 human donor samples (+ 1 pool for normalization) to be condensed into a single tube that could then be deconstructed via high-sensitivity, online, nano liquid-chromatography/mass-spectrometry proteomics. This allowed us to generate a posttranslational molecular map of factors in Scz patient-derived tissue/organoid samples. Consequently, we were able to identify that Scz organoids principally differed from healthy Ctrls due to differences in the total quantity of molecular factors (rather than their diversity), the expression of an ensemble of neuronal factors, and the differential regulation of specific GWAS-implicated [33] disease candidates (namely, PTN and PODXL).

### Convergence upon Depletion of Neuronal Factors in Scz Organoids

The first phenotype to arise in our molecular mapping of Scz organoids was the extent to which canonical neuron identity and development factors were depleted in Scz patient-derived organoids. For several decades, numerous theories have emerged which link neuronal and synaptic function with Scz [36–38], particularly as it relates to cortical dysfunction [39–41] and the cognitive symptoms [42, 43] observed in clinical cases [44]. Recently, progress has been made in understandingly early-arising changes within the developing brain that may influence novel neurodevelopmental factors with putative links to Scz [45]. This has led to numerous investigations of early-arising biological phenomenon in various model systems. Human-derived models, usually leveraging the power of gene edited or patient-derived iPSCs, have consequently revealed alterations in neuronal differentiation [46], mitochondrial metabolic function [47, 48], catecholamine levels [49], neuron-glia interactions [50], synaptogenesis [51], and synaptic function [52]. Thus, patient-derived iPSCs have proven to be a powerful tool in tracing early neurodevelopmental features of Scz [53], which can be further exploited if used to generate human-derived organoids which recapitulate endogenous self-regulatory mechanisms associated with cortical patterning and development within a 3D macroenvironment [11]. Building upon prior Scz organoid work [1, 27, 29, 54], here we report lower levels of an ensemble of neuron-related development factors comprising GAP43, CRABP1, NCAM1, and MYEF2 as well as identity factors comprising MAP2, TUBB3, and SV2A. Broadly speaking, these molecular findings are consistent with our prior work which reported disrupted neurogenesis and lower total neuron numbers within Scz cerebral organoids [1, 55, 56] – a phenotype which has also been independently reported by other groups [28]. Thus, fewer neurons will result in less MAP, TUBB3, and SV2A expression, which is consistent with the molecular outcomes of this independent investigation. Our detection of lower NCAM1 protein levels in Scz organoids is also consistent with a prior report that reported decreased NCAM1 expression in Scz neural progenitor cells [57]. Alterations in the growth-associated factor GAP43 have also been observed across multiple brain regions and independent studies that have evaluated postmortem Scz patient tissue [58–62]. When combined, these data support the idea, and data previously reported in the organoid literature [1, 28], that a depletion in factors supporting neuronal development yields an upstream depletion of neurons within Scz patient-derived organoids [1, 28].

### Regulation of Novel GWAS Factors (PTN & PODXL) in Scz Organoids

The other major phenotype identified in our molecular mapping of Scz cerebral organoids was the differential expression of two novel GWAS factors, namely PTN and PODXL. This analysis comprised us cross-referencing the highest-confident GWAS factors identified in unbiased clinical samples (see [33]) with our complete list of differentially expressed proteins. In our prior report utilizing a smaller 2×2 TMT-LC/MS cohort design [1], we identified the differential expression of four GWAS candidates in Scz cerebral organoids at the protein level (PTN, COMT, PLCL1, and PODXL). Of these candidates, we were able to detect and replicate the differential expression of two of these factors in our much larger sample of *n* = 25 reported here. This specifically comprised alterations in PTN (down-regulated) and PODXL (up-regulated). These factors represent high-confidence GWAS factors associated with Scz, but otherwise have relatively unknown disease relevance. PTN has also been reported to be depleted in neural progenitors and shown to regulate both neurogenesis and survival phenotypes in Scz cerebral organoids [1], providing the first functional molecular data related to this candidate within the Scz literature. Other groups have also recently identified that PTN secreted from neural stem cells supports the maturation of new-born neurons [63], and can function as a neurotrophic growth factor *in vivo* to modulate neuronal loss [64] and long-term potentiation induction [65]. PTN has also since been implicated in a novel amphetamine-model of relevance to Scz [66], a recent computational protein-network analysis underlying Scz [67], as well as at least one nascent Scz gene-association study (*n* = 1,823 humans) [68]. On the other hand, little work has been completed on the role of PODXL in Scz, probably because PODXL is a renal-enriched factor most often associated with kidney podocytes and mesothelial cells [69]. Of note, PODXL has recently been shown to play a role in neurite outgrowth, branching, axonal fasciculation, and synapse number [70], supporting a potential role for this factor in synaptic plasticity. Additionally, PODXL was recently shown to be an apical determinant that may alter lumen size of neural progenitor cell rosettes during morphogenesis [71]. Thus, PODXL may be a fruitful target for future investigations seeking to deconvolute the role of novel Scz GWAS factors within the developing brain.

### Other Novel Differentially Expressed Candidates in Scz Organoids

Lastly, it is worth emphasizing several other differentially expressed molecular candidates observed in Scz cerebral organoids hold biological interest. First and foremost, we identified that Carboxypeptidase E (CPE) was downregulated in Scz cerebral organoids. CPE is a prohormone-processing enzyme [72] and regulated secretory pathway receptor [73], possibly best known for regulating the sorting and activity-dependent secretion of BDNF [74, 75] as well as TrkB surface insertion [76] in neurons. However, CPE was recently suggested to also function as a growth factor independently of its enzymatic and sorting activities [77]. Indeed, amongst other reports suggesting a role in neuroprotection [78], it has recently been shown that CPE regulates cortical neuron migration and dendritic morphology [79]. However, the degree to which these effects is dependent upon its cargo, which includes other growth factors (e.g. BDNF), remains unclear. Lastly, the other notable differentially expressed candidates worthy of discussion comprised alterations within the apolipoprotein family, specifically APOM, APOA1, APOE, APOC3, and APOB. Apolipoproteins have been previously investigated as potential metabolic-related biomarkers [80] in peripherally accessible biological fluids (e.g. CSF [81] or plasma [82]). This specifically includes alterations in APOE and APOA1 in Scz patients [83]. These findings are broadly related to cholesterol [84], fatty acid [85], phospholipid metabolism [86], as well as other membrane-related [87] hypotheses of Scz (which are all somewhat related and/or derived from similar evidence pools). Nonetheless, it is interesting that evidence related to these hypotheses was detectable and reproducible across our sample of patients, and may indicate that further work on potential metabolic factors may also be a further avenue of fruitful research.

### Concluding Remarks

In closing, we identified a broad reduction in molecules important for neuronal identity and development as well as specific alterations in novel GWAS and other disease-relevant molecules previously implicated in Scz. This work collectively supports the idea that Scz is a complex disease underscored by multifaceted changes that likely yield cell-specific as well as multiple mechanisms [55]. In closing, the authors hope that the current dataset may provide insight for other researchers and labs that have an interest in biological data from human-derived 3D stem cell systems but otherwise employ other model systems.

## CONTRIBUTIONS

M.N. and D.C. conceived the project and designed experiments. M.N. generated all 3D tissue from human stem cells, and wrote the manuscript with input and supervision from D.C (senior author). Our technician, A.L., provided important logistical support by assisting with the generation and processing of 3D human-derived tissue. Lastly, H.F. and D.G. completed all LC/MS computational analysis presented in the manuscript, with D.G. serving as the senior author overseeing bioinformatics analyses.

## ACKNOWLEDGEMENTS

M.N. was the recipient of a NHMRC CJ Martin Fellowship that supported mRNA degradation and stem cell training completed at Weill Cornell Medical College of Cornell University.

## METHODS

### Induced Pluripotent Stem Cells

Briefly, human stem cells were principally acquired from NIH deposits at the Rutgers University Cell and DNA Repository. The benefit of utilizing NIH deposited lines is that all biologics have been characterized for identity, pluripotency, exogenous reprogramming factor expression, genetic stability, and viability. In sum, we sampled a total of 25 different iPSC lines comprising both healthy Ctrls and idiopathic Scz patients. Cerebral organoids were generated from all donors in this study, and each iPSC line was biologically independent (representing a unique human donor). Ctrl iPSC lines utilized for cellular experiments included MH0159019, MH0159020, MH0159021, MH0159022, MH0167170, MH0174677, and MH0174686. One Ctrl line (GM23279) was sourced from the Coriell Institute for Medical Research. Scz iPSC lines included MH0159025, MH0159026, MH0185223, MH0185225, MH0200865, MH0217268, MH0185900, MH0185954, MH0185958, MH0185963, MH0185970, MH0185912, MH0185945, MH0185964, MH0185966, MH0185925, and MH0185928. Clinical information for Scz patients is available in Table S1 of our prior publication [1]. All Scz samples were derived from idiopathic cases, which we define here as schizophrenia cases that maintained unknown disease origins and do not meet a genetic/syndrome-based diagnosis (as listed in NIH/NIMH notes). Ctrl iPSC lines were screened for both personal, and family history, of major mental illnesses. All iPSC lines were maintained on Vitronectin-coated plates and fed with Essential 8 (E8) + E8 supplement media (ThermoFisher, CAT#: A1517001).

### 3D Cerebral Organoid Tissue Generation

We adapted the same undirected-differentiation organoid system that we used in our previous, more extensive, analysis of Scz neurodevelopmental mechanisms [1], which had been previously published by Lancaster et al. in *Nature* [17] and *Nature Protocols* [88]. Briefly, 2D iPSC colonies were dissociated and cultured into 3D embryoid bodies in ultra-low attachment plates (Corning; CAT#: 3474). Rock inhibitor (1:1000; Stem Cell Tech, CAT#: 72304) and basic fibroblast growth factor (Pepro Tech, CAT#: 100-18B) are included in media for the first 2-4 days of embryoid body culturing to promote stem cell aggregation and survival. Following this, healthy embryoid bodies are isolated and transferred to Nunclon Sphera 24 well plates (Thermo Scientific, CAT#: 174930) for neural fate specification, using neural induction media. Successful early ‘organoids’ were embedded in a 30μl Matrigel (Corning, CAT#: 354234) spheroid-droplet and polymerized at 37°C for 20-30min which provided a matrix for subsequent neural expansion. Organoids suspended in matrigel droplets were next cultured in terminal organoid media for 4-6 days without agitation, and then cultured with agitation at 60-70RPM until harvested for experiments. For further organoid protocol detail, including QC steps, please refer to our previous publication [1]. Likewise, for further insight into organoid handling for proteomic analysis, please refer to our other organoid manuscript [11].

### Proteomics Sample Preparation, TMT Labeling, & Liquid-Chromatography/Mass-Spectrometry

Isobaric stable isotope labeling was achieved viaTandem Mass Tag pro (TMTpro) chemistry and Liquid-Chromatography/Mass-Spectrometry (LC/MS) proteomics as previously described [1, 11, 66]. Briefly, intact organoids were reduced with dithiotreitol and underwent alkylation with iodoacetamide before tryptic digestion at 37°C overnight. For barcoding chemistry, we employed TMTpro 16-plex labeling according to the manufacturer’s instructions (Thermo Fisher Scientific, CAT# A44521). Each multi-plex experiment contained relevant organoid samples with an additional pooled isobaric reference label made up of the same peptide digest from the pooled mix of organoids (for data normalization between runs; TMT Tag 134N for both TMT-LC/MS runs). A list of sample labeling strategies and replicates is available in the PRIDE proteomics exchange repository. TMT-labelled peptides were desalted using C18’ stage-tips prior to LC-MS analysis. An EASY-nLC 1200, which was coupled to a Fusion Lumos mass spectrometer, (Thermo Fisher Scientific) was utilized in positive, data-dependent acquisition mode, with samples analysed in technical duplicate. Buffer A (0.1% FA in water) and buffer B (0.1% FA in 80% ACN) were used as mobile phases for gradient separation. TMT-labeled peptides were analyzed on a 75 μm I.D. column (ReproSil-Pur C18-AQ, 3μm, Dr. Maisch GmbH, German) was packed in-house. A separation gradient of 5–10% buffer B over 1min, 10%-35% buffer B over 229min, and 35%-100% B over 5min at a flow rate of 300 nL/min was adapted. An Orbitrap mass analyzer acquired Full MS scans over a range of 350-1500 m/z with resolution 120,000 at m/z 200. The top 20 most-abundant precursors were selected with an isolation window of 0.7 Thomsons and fragmented by high-energy collisional dissociation with normalized collision energy of 40. The Orbitrap mass analyzer was also used to acquire MS/MS scans. The automatic gain control target value was 1e6 for full scans and 5e4 for MS/MS scans respectively, and the maximum ion injection time was 54ms for both.

### Data Processing and Bioinformatics Pipeline for Quantitative Analysis

Mass spectra were pre-processed as described [1, 11, 66] and processed using MaxQuant [89] (1.5.5.1). Spectra were searched against the full set of human protein sequences annotated in UniProt (sequence database Sep-2017) using Andromeda. Data was searched as described [1, 11] as a separate and single (combined) batches, with fixed modification, cysteine carbamidomethylation and variable modifications, N-acetylation and methionine oxidation. Searches were performed using a 20 ppm precursor ion tolerance for total protein level analysis. Further modifications included TMT tags on peptide N termini/lysine residues (+229.16293 Da) set as static modifications. Data was processed using trypsin/P as the proteolytic enzyme with up to 2 missed cleavage sites allowed. Peptides less than seven amino acids were not considered for further analysis because of lack of uniqueness, and a 1% False-Discovery Rate (FDR) was used to filter at peptide and protein levels. Protein identification required at least two unique or razor peptides per protein group. Contaminants, and reverse identification were excluded from further data analysis. Quantification was performed with the reporter ion quantification normalization in MaxQuant. Protein intensities were log2 transformed using Perseus [90] (1.×.10). The violin plots of log2 transformed protein intensity distribution and the boxplot of coefficient of variations per sample group were visualized using R package ggplot2. Proteins quantified in at least 70% of samples in at least one sample group were subjected to downstream visualization (principal component analysis, volcano plot) and statistical analysis using Perseus. For principal component analysis, missing values were imputed from normal distribution (downshift 1.8, width 0.3) using Perseus. For differential expression analysis proteins were subjected to Welch’s t-test; p-value < 0.05 and |log2FC|>0.5 visualized in volcano plot and subjected to downstream functional enrichment analysis using g:Profiler, including Gene Ontology, KEGG and Reactome databases (as described, [91, 92]).

### Data Availability Statement

The MS proteomics raw data and MaxQuant search parameters have been deposited to the ProteomeXchange Consortium (http://www.proteomexchange.org/) via the PRIDE partner repository [93] with the data set identifier PXD027812.

